# WINDEX: A hierarchical integration of site- and window-based statistics for characterizing the footprint of positive selection in genome-wide population genetic data

**DOI:** 10.64898/2026.03.26.714384

**Authors:** Hannah Snell, Scott McCallum, Dhruv Raghavan, Ritambhara Singh, Sohini Ramachandran, Lauren Sugden

## Abstract

Adaptive mutations, or mutations that confer a fitness benefit, can leave behind distinct signals in genetic data. Computational methods have improved the localization of adaptive mutations in genetic samples using a range of statistical and machine learning classification techniques. However, these methods miss the opportunity to jointly integrate statistics at both the site and window-based level, thus failing to harness all available statistical evidence to detect selection. Our method, WINDEX, combines these different resolutions of statistics to improve the detection of adaptive mutations among hitchhiking signals. Our model simultaneously integrates emissions at different resolutions by defining site-based and window-based latent states corresponding to neutral, linked, and sweep regions, with the site-based states and transition models nested within the window-based states. Using evolutionary simulations with varying selection parameters, we validate the ability of WINDEX to classify positive selective sweeps. Using data from the 1000 Genomes Project, we show that WINDEX is able to identify regions harboring signals of selective sweeps, and provides improved localization within those regions over existing methods. In addition, using WINDEX genome-wide allows for estimation of the proportion of whole genomes that are under positive selective pressures; our estimates of between 9.7-10.5% across different populations provide support for other preliminary estimates of these quantities.

**Author summary:** Population geneticists often seek evidence for positive selective sweeps, or an evolutionary event in which a beneficial allele increases in frequency over time in a population, resulting in increased fitness of the individuals that have said allele. Positive selective sweeps, however, are difficult to detect due to varying patterns of linkage disequilibrium (LD), or the nonrandom association of alleles, and detecting these signals reliably among differing LD structures remains a challenge in the field. In this work, we present WINDEX, a probabilistic framework designed to leverage signals of positive selective sweeps at both the site- and window-levels in the form of a hierarchical hidden Markov model (HHMM), to localize regions of positive selective sweeps in aligned haplotype data. We validate WINDEX in evolutionary simulations over varying positive selective sweep scenarios, showcasing the improved resolution that the HHMM structure provides. We apply WINDEX in comparative genomic scans of canonical sites of positive selection as well as whole-genome scans to demonstrate the tool’s power in localizing functionally-validated signals of selection and to offer insights into the proportion of the human genome currently under positive selective pressures. WINDEX is publicly available and easy to apply to many cases of human genetic data.

## Introduction

A central focus in population genetics is the development and application of statistical approaches to identify strongly adaptive mutations in genetic data. Strongly adaptive mutations can rise quickly in allele frequency in populations via processes called positive selective sweeps. A positive selective sweep can originate from a *de novo* beneficial mutation being introduced into a population (known as a “hard sweep”) or arise from standing variation, a preexisting mutation that becomes advantageous in a population [1]. Traditionally, scans for mutations undergoing positive selective sweeps are conducted by calculating one or more statistics on population genetic data that detect different identifying signatures of positive selection. Many statistics can be calculated at the individual site level. These typically measure changes in allele frequencies, as in the case of the Fixation Index (*F*_*ST*_) [2], or the frequencies of shared haplotypes, as in the case of the Integrated Haplotype Score (iHS) [3]. Window-level statistics such as Tajima’s D [4], measure deviations from neutrality of the site frequency spectrum over a user-specified window size. In a positive selective sweep, it is common for the selected allele and nearby “hitchhiking” alleles within a recombination window to both increase in frequency. This correlation in genotype states, as measured by linkage disequilibrium (LD), poses challenges in the localization of selected alleles in positive selection scans. In addition, LD patterns vary depending on the demographic history of the individuals studied, which is caused by different histories of population migration and expansion that affect recombination patterns [5–11]. Also outlined in supplementary figures 17-19 of Sugden *et al*., 2018, a challenge that positive selection scanners such as SWIF(r) face when classifying candidate sweep regions is variation in the frequency of sweep signals based on population, which results from confounding by differing lengths of LD blocks and recombination rates between populations [12–14]. The selection statistics used in scans for characterizing adaptive mutations can be sensitive to this LD variation, therefore making a classifier less generalizable depending on the observed population structure, and this remains underexplored as an area for improvement in positive selection classification.

In addition to univariate measures to detect selection, population geneticists have developed computational methods combining multiple selection statistics to identify adaptive mutations in population genetic data. These methods vary in model type and complexity: for example, iSAFE [15] is a scoring method in which alleles are ranked based on their contribution to the overall selection signal. Alternatively, S/HIC [16] is an Extra-Trees classifier that makes multilevel classifications (“neutral”,”linked”,”sweep”) using window-based statistics and was developed to delineate hard versus soft selective sweeps. Finally, SWIF(r) [17], is a probabilistic framework that uses averaged one-dependence estimators to make binary classifications (neutral vs. sweep mutations) from population genetic haplotype data, and offers more input statistic flexibility as long as they are all at the same resolution level in the data (site or window). However, all of these methods only use site-based or window-based statistics, not both simultaneously. Because these different statistical scales are sensitive to different timescales of positive selective sweeps (for example, the site-frequency spectrum, which can be analyzed by the use of both site- and window-based statistics, helps to identify older sweeps while long shared haplotypes are sensitive to recent sweeps [9]), existing methods miss out on integrating both scales of selection statistics to inform classifications.

To address these challenges, we introduce WINDEX: a framework to characterize the footprint of positive selection on genome-wide data using a hierarchical hidden Markov model (HHMM) that integrates site-based and window-based summary statistics to make classifications at both statistical resolutions. We generated evolutionary simulations representing varying conditions of positive selective sweeps in three populations to train, test, and validate WINDEX against its predecessor tool, SWIF(r) [17]. Using data from the 1000 Genomes Project [18], we compared the localization ability of WINDEX at the site-level against iSAFE [15] using canonical sites of positive selection and applied WINDEX as a genome-wide scanner to estimate the proportion of the human genome currently under positive selective pressures, comparing these results to those from a genomic scan using S/HIC [16]. WINDEX shows comparable localization to iSAFE, and with the use of stochastic backtrace at the site-level, shows a reliable uncertainty measurement for localizing sweep signals. At the whole-genome level, WINDEX offers additional evidence for better understanding genome-wide positive selective sweep patterns. WINDEX is an interpretable probabilistic framework that improves localization of adaptive mutations and understanding of the influence of LD in positive selective sweeps.

## Materials and methods

### Overview of the WINDEX model

WINDEX is a hierarchical hidden Markov model (HHMM) that incorporates two main state levels: a window-based level, whose emissions are window-based statistics (e.g. Tajima’s D [4]) that are calculated genome-wide in user-defined window sizes, and a site-based level, whose emissions are site-based statistics (e.g. iHS [3]) that are calculated per-site in genome-wide data (Fig 1A). These levels are nested such that each window state gives rise to a sequence of site states, with the allowable site states and transitions dependent on the window state. WINDEX contains three main types of states: “neutral” states, which correspond to neutrally evolving windows or sites, “linked” states (defined as “linked left” and “linked right” where the use of “left” and “right” allow us to algorithmically enforce a single sweep to be localized), which correspond to windows or sites considered linked or hitchhiking with a positive selective sweep, and “sweep” states, which correspond to, at the window-level, windows that contain a sweep site, or at the site-level, individual sweep sites. Both levels of emissions in WINDEX are calculated using a Naïve-Bayes framework where the product of each statistical value per class is reflected. A full state path through this model begins in a window state, then moves through site states for each site contained in that window, before returning to the window level and transitioning to the next window state. This particular design allows for each statistical scale to inform the other. In order to achieve high probability, a path through the model must attend to the emission and transition probabilities at both levels simultaneously. Using a hierarchical extension of the Viterbi algorithm [19], WINDEX reports the window- and site-based paths that correspond with the maximum likelihood path through the model, outputting separate site- and window-based files (Fig 1B).

**Fig 1.**
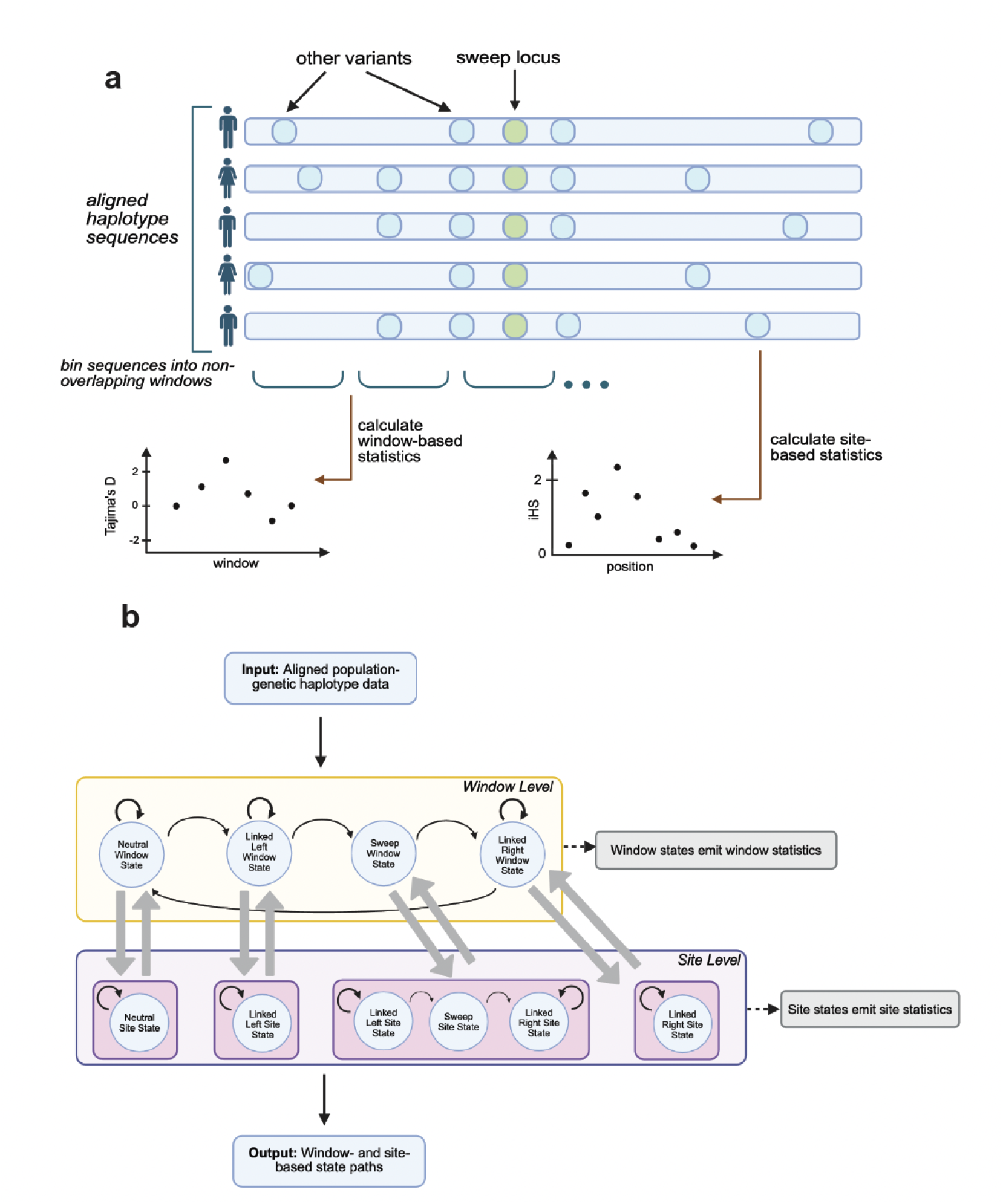
Method outline and state structure of WINDEX. (Fig. 1A) WINDEX requires input of summary statistics calculated from aligned haplotype data, both at the non-overlapping window level (such as Tajima’s D [4]) and the site-level (such as iHS [3]). (Fig. 1B) WINDEX is a hierarchical hidden Markov model (HHMM) consisting of two main state levels: a window-based level that operates at a user-defined window size resolution with window statistics as emissions, and a site-based level that operates at an individual variant resolution with site statistics as emissions. WINDEX works by initializing first at the window level, then moving through the sites within that window in one of the site-level substructures, then transitioning back to the window level for the next window state transition. The window-level transitions are controlled by a single transition matrix, while the site substructures each have their own transition matrices. See Methods for emission training description and Supplementary Tables S3 and S4 for transition probability definitions.

The WINDEX site-level state structure is completely dependent on the window-level states, thereby restricting the allowable site states at different windows in the model. More specifically, when the model transitions into a neutral window state, the sites within this window can only transition into neutral site states. This is also true of both linked window states, where both only give rise to linked site states. In a sweep window, the model can transition at the site level from linked “left” of the sweep site, to the sweep site, then linked “right” of the sweep site.

### Evolutionary simulations

To train and test WINDEX, we generated genetic simulations using the Three Population Out of Africa demographic model [20] with SLiM v4.0 [21]. We simulated a single hard *de novo* sweep in the center of a contig of size 1 million base pairs for each population with varying sweep scenarios described in Table 1. We also generated corresponding neutral simulations for each population with the same demographic parameters but without selection. In total, our simulation set contained 1500 sweep simulations and 300 neutral simulations (100 simulations per population-scenario combination).

**Table 1.** Simulation conditions for training WINDEX.

| Mutation Introduction Time | Selection Coefficient | Average FAF | Num Simulations $\geq 60\%$FAF |
| --- | --- | --- | --- |
| 300 generations | 0.05 | 84% | 232 |
| 300 generations | 0.02 | 34% | 0 |
| 300 generations | 0.01 | 32% | 0 |
| 600 generations | 0.02 | 75% | 196 |
| 600 generations | 0.01 | 41% | 15 |

Using SLiM [21] and following the Three Population out of Africa [20] human demographic model, we introducted a single *de novo* hard sweep mutation according to the above table parameters in each of the three model populations (YRI, CEU, and CHB). Since SLiM is a forward simulator, the average final allele frequency (FAF) of the sweep mutation was estimated across all training simulations (column 3). A subset of these simulations were then used based on their sweep mutation final allele frequency (set *≥* 60%) to ensure that only simulations with strong selection would contribute to training and testing WINDEX (column 4). See Supplementary Figure S1 for population-level sweep site final allele frequency distributions in simulations. See WINDEX Github for example simulation code.

### Calculation of input statistics

WINDEX requires two statistic files per aligned genetic sequence input: one site-based statistic file and one window-based statistic file. Additionally, to train the emission matrix, WINDEX requires an input of the marginal distributions of each site- and window-based statistic. An advantage of WINDEX is that it flexibly takes as input any statistic that a user provides, as long as there is at least one window-based statistic and one site-based statistic calculated for all user-defined windows and corresponding sites in the data. For the validation and application of WINDEX in this study, we chose a diverse set of site- and window-based statistics (Supplementary Table S1 and S2). We chose to use a 40-kilobase size window for all window-based statistics in our simulations, centering the sweep signal within the middle 40kb window on the contig, resulting in 25 equally sized contigs. After computing all statistic values across all simulated contigs, we calculated the global neutral average and global neutral standard deviations within each population of each statistic and used these to standardize all neutral and sweep files. For benchmarking, we split the neutral and sweep simulations into training and testing data in an 80:20 ratio, respectively.

#### Site-based statistics

We used selscan v.2.0 [22] to calculate three site-based statistics: Integrated Haplotype Score (iHS) [3], Number of Segregating Sites by Length (nSL) [23], and Cross-Population Extended Haplotype Homozygosity (XP-EHH) [1]. Additionally, we implemented VCFtools v1.14 [24] to calculate Fixation Index (*F*_*ST*_) [2]. Finally, we calculated the change in derived allele frequency (ΔDAF) of each SNP using the ‘vcftools –freq’ command to produce derived allele frequencies of each allele and subtracted the average derived allele frequency of the two non-focal populations (see Github). When performing standardization of these statistics, we used selscan’s ‘norm’ tool to standardize values of iHS, nSL, and XP-EHH to the global neutral average in allele frequency bins of 20 base pairs. Supplementary Table S1 for tabular summary of these statistics.

#### Window-based statistics

We applied scikit-allel v.1.3.3 [25] to calculate five window-based statistics: Π [26], Watterson’s *θ* [27], and Tajima’s D [4], Garud’s H [28], and Population Branching Statistic [29]. We also calculated three additional statistics: Fay and Wu’s H [30] was calculated using the Π estimator, Zeng’s E [31] was calculated using the Watterson’s *θ* estimator, and NSS (number of segregating sites per window) was calculated by using the scikit-allel function “is_segregating()”. All window-based statistics were standardized against the global neutral average values of each statistic before split into the same training and testing files as the site-based statistics at an 80:20 ratio. See Supplementary Table S2 for tabular summary of these statistics.

### Architecture and evaluation of WINDEX

Our methodological goal is for a user to be able to leverage statistical evidence from both site- and window-based statistics to detect regions undergoing a selective sweep in haplotype data. The crux of developing WINDEX is integrating emissions from both site- and window-based statistics, even though they operate at two different resolutions in the data. To do this, we constructed WINDEX with two levels of states, one at the window-based level, and one at the site-based level, where the state path of the window level informs the site state substructure in which the model continues (1B). This structure is inspired by the multi-timestep hierarchical hidden Markov model described in Adam *et al*. (2019) for aquatic creature movement and feeding patterns [32]. This paper applied an HHMM with emissions at multiple levels, which is not the typical use-case for HHMMs. WINDEX is formally defined as:

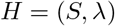

where S is the set of all possible hidden states:

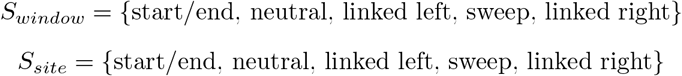

and *λ* represents the site- and window-based transition and emission probabilities:

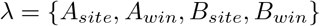

In WINDEX, *S* contains five unique states at both emission levels. The linked regions are partitioned into two *linked* states, labeled based on their relative positions to the sweep state location (i.e.,”left of the sweep location” or “right of the sweep location”). This forces the model to go through the sweep state on either level controlled by the transition probabilities, therefore guaranteeing a sweep classification and avoiding linked regions that are not associated with sweeps. The *start/end* state captures the initial and terminal probabilities of each substructure of the model that a sequence iterates through, so the initial probabilities and terminal probabilities at any step in the framework are defined in the transitions *A*, where *A*_*window*_ controls transitions for the window level of the model, and three separate *A*_*site*_ matrices control the site-level transitions. WINDEX uses different site state structures and transition probabilities (*A*) depending on the state sequence of the window-based level. Additionally, WINDEX has emissions (*B*) at both levels to account for observations made from both window- and site-based statistics. The following is the formal definition of the WINDEX model.

First, we define a site-based path through the model within window *t*, assuming window *t* is in window state *r*:

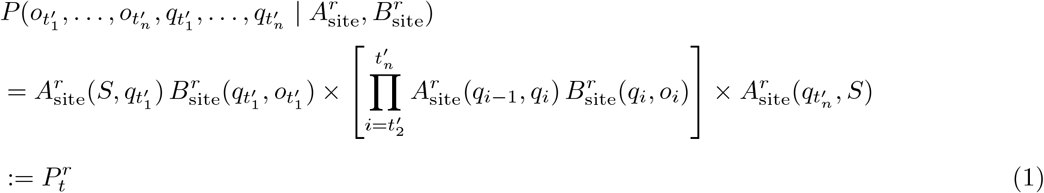

Then, using this definition, we define a hierarchical window-based path through the model:

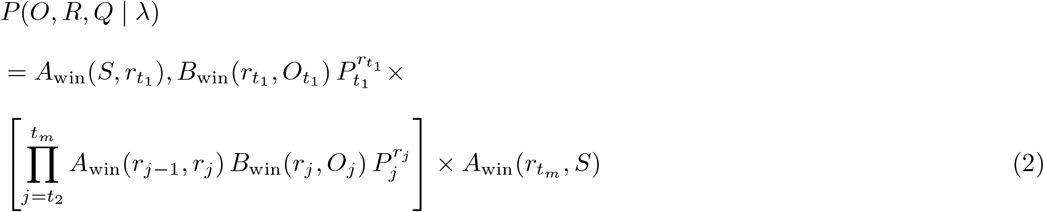

where

*O* is the sequence of observations (uppercase for window, lowercase for site, *R* is the sequence of window states,

*Q* is the sequence of site states,

*A* is the state transition matrix,

*B* is the emission matrix,

*r* indexes the window state, *q* indexes the site state,

*t* denotes the window-state time step, *t*^*′*^ denotes the site-state time step.

*S* denotes the fixed start/end state

*λ* denotes the set of transition and emission probabilities *A*_win_, *B*_win_, *A*_site_, *B*_site_

WINDEX uses emissions that are specific to each level. For both levels of emissions, we employed a Naïve Bayes likelihood function to combine all the evidence for each statistic probabilistically. To do this, we generated marginal distributions of each normalized statistic from the simulation training data. We then computed the product of these marginal distributions so that the model had a representation of the probability of emitting an observation, given both the hidden state and all information provided by the input statistics. WINDEX has five main transition probability matrices: one for the window-level, and four for the site level. This ensures that, for each window state, there is a deterministic set of site states that are controlled by independent site-level transition matrices. Given this structure, WINDEX is highly flexible, but also has many transition parameters that must be tuned. Because of this, we have parameterized all of these transitions to provide a set of “default” transitions for a user to begin training with WINDEX. These parameterizations are controlled by two main biologically-relevant features of a user’s training haplotype data: the expected size of a linked region surrounding a hard sweep, and the expected frequency of hard sweep loci in the training data (Supplementary Table S3). These parameterized transitions can then be further tuned to best fit the particular biological application. The manually tuned probabilities used for all of our runs of WINDEX results in this paper are reported in Supplementary Table S4. For full transition matrix implementation, please see the WINDEX Github.

We evaluated WINDEX using an extended version of the Viterbi algorithm (Viterbi, 1967) that incorporates the hierarchical structure. To do this, we performed a full forward pass through the hierarchical structure and obtained the most likely window state conditional on the previous window state and site-based state sequence. Then, WINDEX calculates a regular Viterbi path for the sites within a site state substructure, which is determined by the window state. The probabilities at the site-level are incorporated into the window-level probabilities so that each window transition is also informed by the site-based data. The algorithm terminates when the window-level Viterbi algorithm reaches the *start/end* state, at the end of the haplotype sequence.

Additionally, we introduced a site-level stochastic backtrace algorithm in our iSAFE comparative analysis results section. This algorithm provides a statistical ensemble of state predictions by first completing a forward pass through the state structure, then completing a backward state assignment where each transition is determined by a weighted probability state vector that is computed based on the final forward probabilities [33]. For complete algorithm code, see the WINDEX Github.

### Real-world data analysis with 1000 Genomes Project

We used variant information from Phase 3 of the 1000 Genomes Project over three populations: Yoruba Nigerians (YRI, 108 genomes of unrelated individuals), Central Europeans of Utah (CEU, 99 genomes of unrelated indivduals), and Han Chinese in Beijing, China (CHB, 103 genomes of unrelated individuals) [18]. We obtained VCF files containing variants for all three populations per chromosome, and performed cleaning steps for our downstream analysis. First, we polarized these files so that an additional INFO column denoted ancestral versus derived allele states according to the Homo sapiens ancestral allele sequence from Ensembl (release 74; ftp://ftp.ensembl.org/pub/release-74/fasta/ancestral_alleles/). We only included variants that had an ancestral or derived state label in our analysis. Next, we filtered each VCF file so that only variants with a minor allele frequency (MAF) *≥* 0.05 would be included. Next we filtered variants that had more than two alleles, so only biallelic SNPs remained. Once the filtering steps were complete, we split each chromosome-level VCF file by population and calculated the same site-based and window-based statistics according to the steps explained for the simulations. For standardization, all statistics were instead standardized against their chromosomal mean and standard deviation.

We ran iSAFE v.1.1.1 [15] in 300kb regions each around two canonical site of positive selection, positioning the locus of interest in the middle 100kb window. We chose this region size to satisfy the minimum scanning region suggested by the iSAFE documentation, while minimizing the region that WINDEX scans over for high localization probability. We used the ‘–MaxRank’ flag to ensure we examined all SNPs in the 300kb region and the ‘–IgnoreGaps’ flag to ignore large gaps between variants.

We ran WINDEX in the same 100kb windows published by Schrider and Kern in 2018 in YRI and CEU genomes [16]. For easy comparison to the S/HIC results produced in Phase 1 1000 Genomes data, we releveled the S/HIC outputs to best match the outputs of WINDEX, since WINDEX is not designed to delineate between soft and hard sweeps. To obtain the proportions reported in Table 1, we summed the number “linkedSoft” and “linkedHard” regions in the S/HIC output to make a single “linked” classification. We did this same releveling for the hard and soft sweep classes: “hard” and “soft” were combined to “sweep”.

## Results

### Performance of WINDEX in evolutionary simulations

We validated WINDEX using a set of evolutionary simulations generated according to the Three Population out of Africa demographic model [20] with positive selection using the scenarios described in Table 1 (see Methods). We trained WINDEX on all simulations that had a final allele frequency of the selected allele above 60% to ensure that the sweeps had sufficiently spread through the target population (Supplementary Fig. S1). The simulations that satisfied this condition overwhelmingly had the parameters 300 generations, 0.05 selection strength and 600 generations, 0.02 selection strength, so we evaluated the performance of WINDEX using the held out testing sets from both of these scenarios. The emission model for WINDEX shares its structure with the Naïve-Bayes version of its predecessor tool, SWIF(r) (we denote this as *NB-SWIF(r)*) [17], so for our first comparison of performance, we ran NB-SWIF(r) at both the window- and site-based levels using the same training and testing simulations as for WINDEX. This comparison allows us to ascertain the advantage of the HHMM architecture against a simpler model while using the same statistics as input. In Figs 2A and 2B, WINDEX performs well at identifying the correct window that contains the sweep mutation (83.3% and 76.7% for 2A and 2B, respectively). Additionally, the windows adjacent to the sweep window are consistently classified as linked windows and the number of testing simulations supporting this classification decreases as one moves further away from the center of the simulated contig. This pattern is consistent with what is expected in a positive selective sweep scenario and demonstrates that, in a strong selection case, WINDEX performs well at localizing sweep windows correctly. When we compare this result to NB-SWIF(r) in Figs 2C and 2D, we see that NB-SWIF(r) shows a similar decay of linked window calls with distance from the true sweep window, but calls the sweep window correctly in a low percentage of testing simulations (both 26.7% for the scenarios in Figs 2C and 2D). These results demonstrate the advantage of the hierarchical framework in WINDEX: window-based NB-SWIF(r) has challenges with localization, but when using a hierarchical model, the window-level classifications are able to leverage the information provided by the site-level statistics in the model as well as information from neighboring windows to accurately localize the true sweep window.

**Fig 2.**
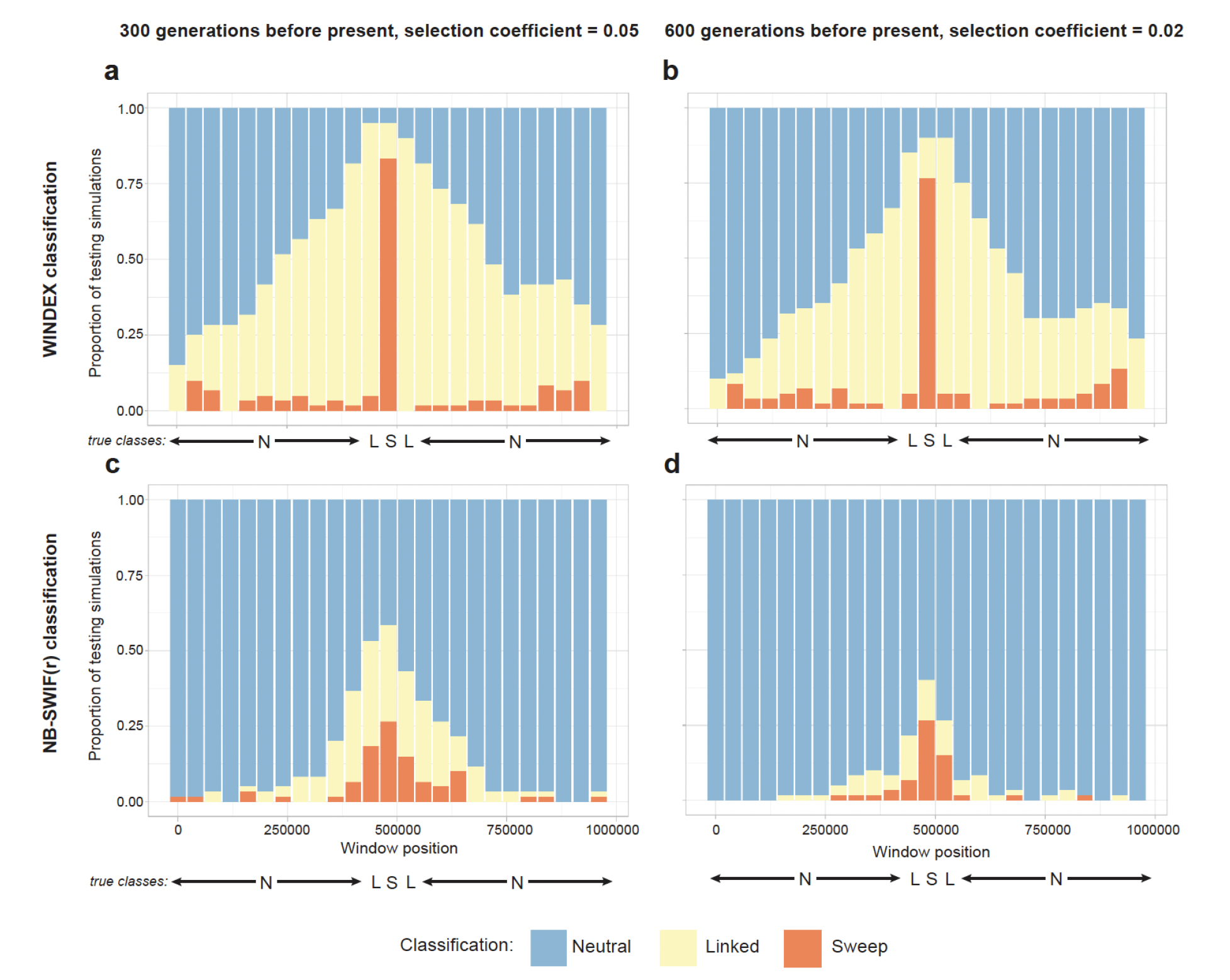
Comparative performance of WINDEX vs. NB-SWIF(r) in evolutionary simulations at the non-overlapping window scale shows superior localization of sweep-containing windows with WINDEX. We trained WINDEX with evolutionary simulations using the Three Population out of Africa demographic model [20] with varying positive selective sweep scenarios (see Table 1). WINDEX was tested in two conditions: 300 generations before present, selection coefficient = 0.05 and 600 generations before present, selection coefficient = 0.02, both based on sweep mutation final allele frequency (*≥* 60%, see **??**). (Figs. 2A and 2B) WINDEX correctly classifies the true sweep window in a large proportion of testing simulations across two strong selection scenarios. The red centered bar shows the number of testing simulations where WINDEX accurately classifies the sweep window. Surrounding yellow bars also show that WINDEX classifies nearby windows as linked with increasingly higher proportions closer to the true sweep window, which is expected in a true positive selective sweep. (Figs. 2C and 2D) The Naive-Bayes version of SWIF(r) (NB-SWIF(r)) [17] was trained with the same simulation set for method comparison. While NB-SWIF(r) has the highest sweep proportion in the true sweep window in both scenarios, the number of testing simulations supporting this classification are much lower than with WINDEX. Similarly, the linked footprint around the true sweep window is diminished compared to the WINDEX classifications (yellow bars).

We made similar comparisons between models at the site level. Fig. 3 identifies strong selection signals at the site level across both WINDEX and NB-SWIF(r), viewed in a 2kb window centered around the true sweep site. In both scenarios, WINDEX demonstrates high probability in identifying the correct sweep site locus within a correctly classified sweep window (94% and 80%, Figs 3A and 3B respectively). NB-SWIF(r) shows better accuracy at identifying the sweep site (100% and 92% Figs 3C and 3D respectively), but also shows a substantially lower specificity since nearby signals in this region are classified as sweeps with similar probabilities in both scenarios, while WINDEX classifies all nearby sites in this window as linked (for full precision and recall metrics, see Supplementary Table S1). Because of the hierarchical structure of WINDEX, sites in a sweep window are restricted to either linked or sweep site states, meaning that sites close to a true sweep locus are likely to be called as linked rather than neutral. Furthermore, the site transition matrix within a sweep window enforces prioritization of the most compelling sweep site, resulting in the increased specificity of sweep site calls (see Methods).

**Fig 3.**
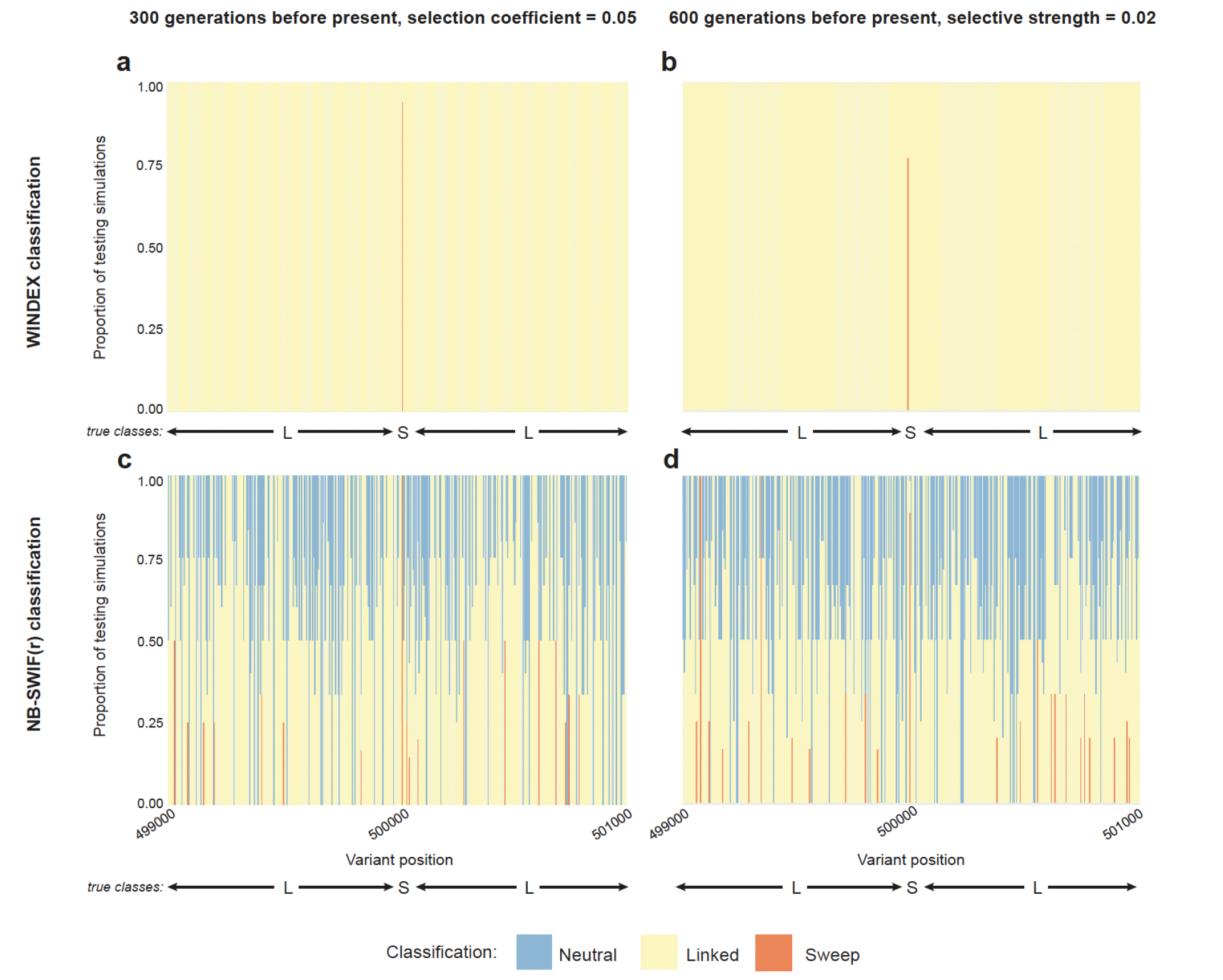
Comparative performance of WINDEX vs. NB-SWIF(r) in evolutionary simulations at the site scale show superior localization of sweep loci using WINDEX. Similar to Fig. 2, WINDEX and NB-SWIF(r) were both evaluated in their abilities to localize the footprint of two strong positive selection scenarios using evolutionary simulations (see Methods). All four panels depict +/-1kb around the true sweep locus inside the true sweep windows denoted by the labels in Fig. 2. (Fig. 3A and 3B) WINDEX classifies the true sweep locus in high proportions of testing simulations (red bars). Additionally, WINDEX accurately classifies sites labeled as linked that are close to the true sweep locus (yellow bars) with no classifications reported for neutral signals. (Fig. 3C and 3D) NB-SWIF(r) also identifies the true sweep locus in high proportions of testing simulations, but struggles to localize this signal within a small genetic window of the true sweep locus. In both scenarios, there are putative sweep sites nearby classified with varying proportions of testing simulations supporting them as well as many true-labeled linked sites classified as neutrally evolving.

### Performance comparison of WINDEX and iSAFE with applications to canonical sites of positive selection

To better understand the ability of WINDEX to detect positive selective sweep footprints in human genomes, we compared site-based classifications from WINDEX against the “integrated selection of allele favored by evolution” (iSAFE) statistic [15]. When given population haplotype information, iSAFE outputs a ranked score for the likelihood that an allele contributes to a positive selection signal. We chose two canonical sites of positive selection to compare the performance of both WINDEX and iSAFE for localizing the validated adaptive mutation: in East Asian individuals, the derived allele (370A) of the rs3827760 variant within exon 12 of the Ectodysplasin A Receptor (*EDAR*) gene that is responsible for the development of hair follicles, teeth, and sweat glands [34], and in European individuals, the derived allele (111A) of the rs1426654 variant of the SLC24A5 gene that is involved in pigmentation of hair, skin, and eye color [35; 36].

Using Han Chinese from Beijing, China (CHB) and Utah residents with Northern and Western European Ancestry (CEU) genomes from the 1000 Genomes Project [18], we compared WINDEX and iSAFE in 300kb regions around the known locus of each canonical variant position (Fig. 4). In Figs. 4A and 4D, we show that iSAFE ranks many candidate signals in the two respective regions, scoring the rs3827760 variant in *EDAR* in the top three signals, and the rs1426654 variant in *SLC24A5* in the top five signals. In Figs. 4B and 4E, we show that WINDEX classifies the selected variants as the only sweep sites in each respective region and surrounding sites as linked sites.

**Fig 4.**
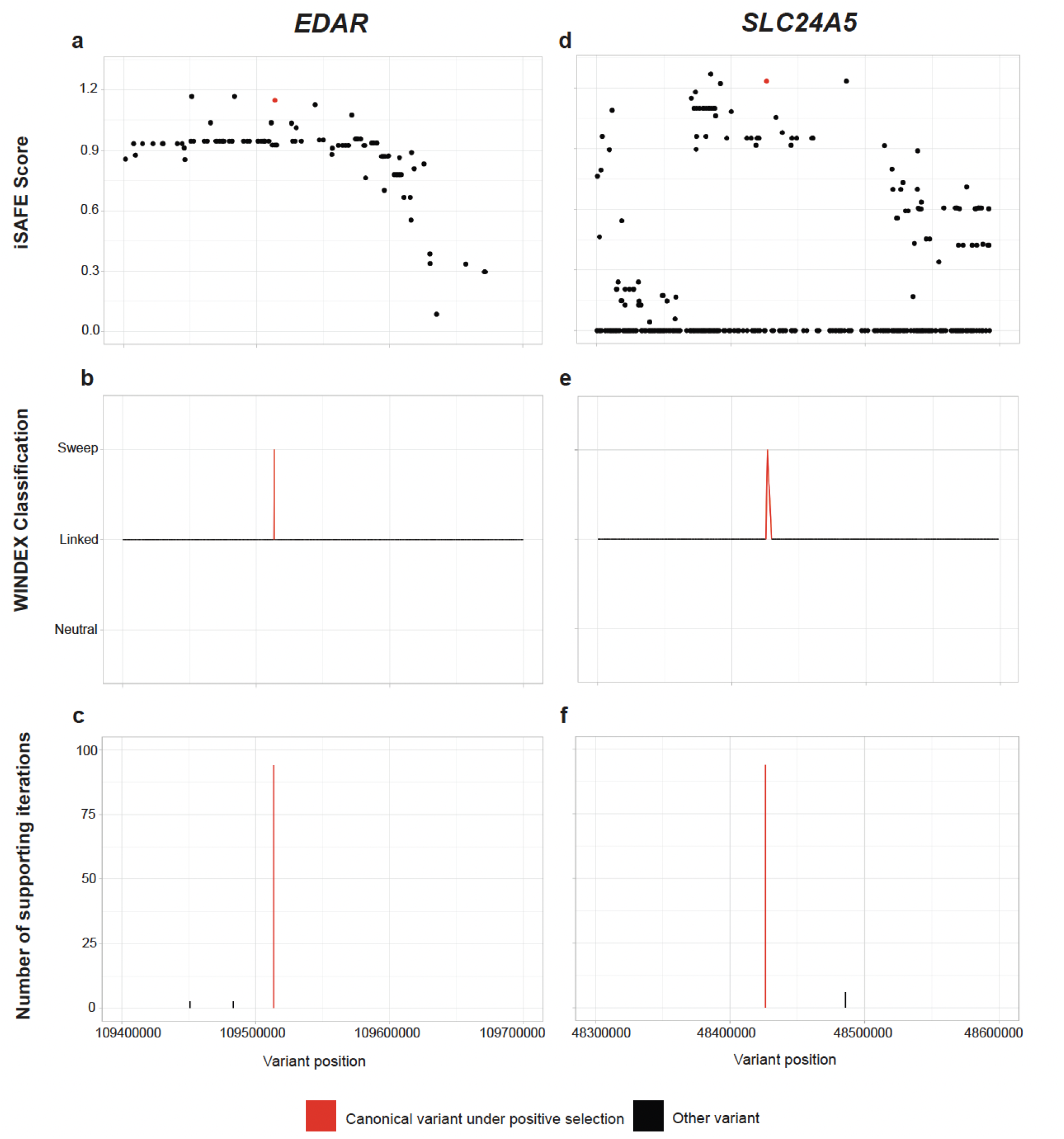
Comparative scans in *EDAR* and *SLC24A5* with iSAFE and WINDEX show that WINDEX outperforms iSAFE in localization of candidate signals of positive selection. Using both iSAFE and WINDEX, 300kb regions of canonical selection signals were scanned in YRI and CEU genomes from the 1000 Genomes Project [18] and compared based on accuracy of localization. (Figs. 4A and 4D) In both canonical regions of positive selection, iSAFE ranks the putative variant within the top five strongest signals, but shows other highly ranked variants close by, which pose localization challenges. (Figs. 4B and 4E) WINDEX accurately localizes the putative variants in both canonical regions of positive selection while classifying all surrounding sites as linked. (Figs. 4C and 4F) Implementation of a stochastic backtrace evaluation algorithm at the site level allows for a measure of uncertainty to be assigned to WINDEX classifications. Over 100 stochastic backtraces, both putative signals are supported 94%, while low-level support is shown for nearby signals.

WINDEX is constrained to identifying only one sweep site classification in its HHMM maximum likelihood path, but an additional strength to using an HMM-based model is the opportunity to create a measure of uncertainty around the maximum a posteriori sweep site via stochastic backtrace. While iSAFE ranks the functionally validated variant high in both *EDAR* and *SLC24A5*, there are many highly ranked variants nearby that are similarly scored, posing a challenge for localization of the selected signal. We applied a stochastic backtrace site-based algorithm to create an uncertainty metric around each WINDEX sweep site classification in the window classified as sweep (Figs. 4C and 4F). We observed that WINDEX combined with 100 stochastic backtrace replicates classifies the selected variants as sweeps with high confidence (94/100 supporting stochastic backtrace iterations in both cases), and also provides some lower support for nearby competing signals classified as sweep sites. The combination of running WINDEX in larger genomic regions plus the use of the stochastic backtrace algorithm for uncertainty measurements at the site-level offers targeted prioritization of candidate adaptive variants at a resolution beyond competing approaches.

### Performance comparison of WINDEX and S/HIC with applications to whole-genome estimations of the proportion of the human genome under positive selection

One motivating application for developing WINDEX is to estimate the proportion of whole-genome samples under the influence of positive selection. S/HIC (Schrider and Kern, 2016) is another selection scanning tool that addressed this question by reporting hard and soft sweep regions classified in whole-genome scans of individuals from 1000 Genomes Project data. To compare results from these two approaches, we ran WINDEX in the same 100kb windows over whole genomes of individuals from 1000 Genomes Project CEU and YRI populations as done with S/HIC for whole-genome classification (see Methods). We then compared the window-level WINDEX classifications to the classifications published with S/HIC [16] in the same populations. Table 2 summarize these whole-genome results. In both CEU and YRI genomes, all three classes show very similar proportions across tools. In CEU, sweep proportions are 0.105 for S/HIC versus 0.097 for WINDEX, linked proportions are 0.5485 for S/HIC versus 0.417 for WINDEX, and neutral proportions are 0.3465 for S/HIC versus 0.486 for WINDEX. Similarly in YRI, sweep proportions are 0.082 for S/HIC versus 0.105 for WINDEX, linked proportions are 0.639 for S/HIC versus 0.496 for WINDEX, and neutral proportions are 0.279 for S/HIC versus 0.399 for WINDEX. Across both sets of population genomes outlined in Table 2, WINDEX consistently classifies proportions of CEU and YRI genomes as sweep regions similarly to S/HIC, and consistently classifies less of these genomes as linked compared to S/HIC. To better interpret these classification results, we examined how many of these classifications overlap between tools at (1) an exact window position, and (2) WINDEX classifications matching within +/-500kb of S/HIC classifications. In both populations, there is a low classification concordance across exact matching windows: In CEU, sweep class colocalization is 15.1%, linked class colocalization is 58.6%, and neutral class colocalization is 37.6%, and in YRI, sweep class colocalization is 9.4%, linked class colocalization is 65.3%, and neutral class colocalization is 30.0%. When examining colocalization within +/-500kb of the S/HIC classifications, we observed an increase in the colocalization across all classes. In CEU, sweep class colocalization is 64.8%, linked class colocalization is 96.2%, and neutral class colocalization is 82.2%, and in YRI, sweep class colocalization is 56.0%, linked class colocalization is 96.6%, and neutral class colocalization is 73.6%. These results are an average across all chromosomes and genomes in each population. To explore per-chromosome results, see the Supplementary Tables S5-S8.

**Table 2.**
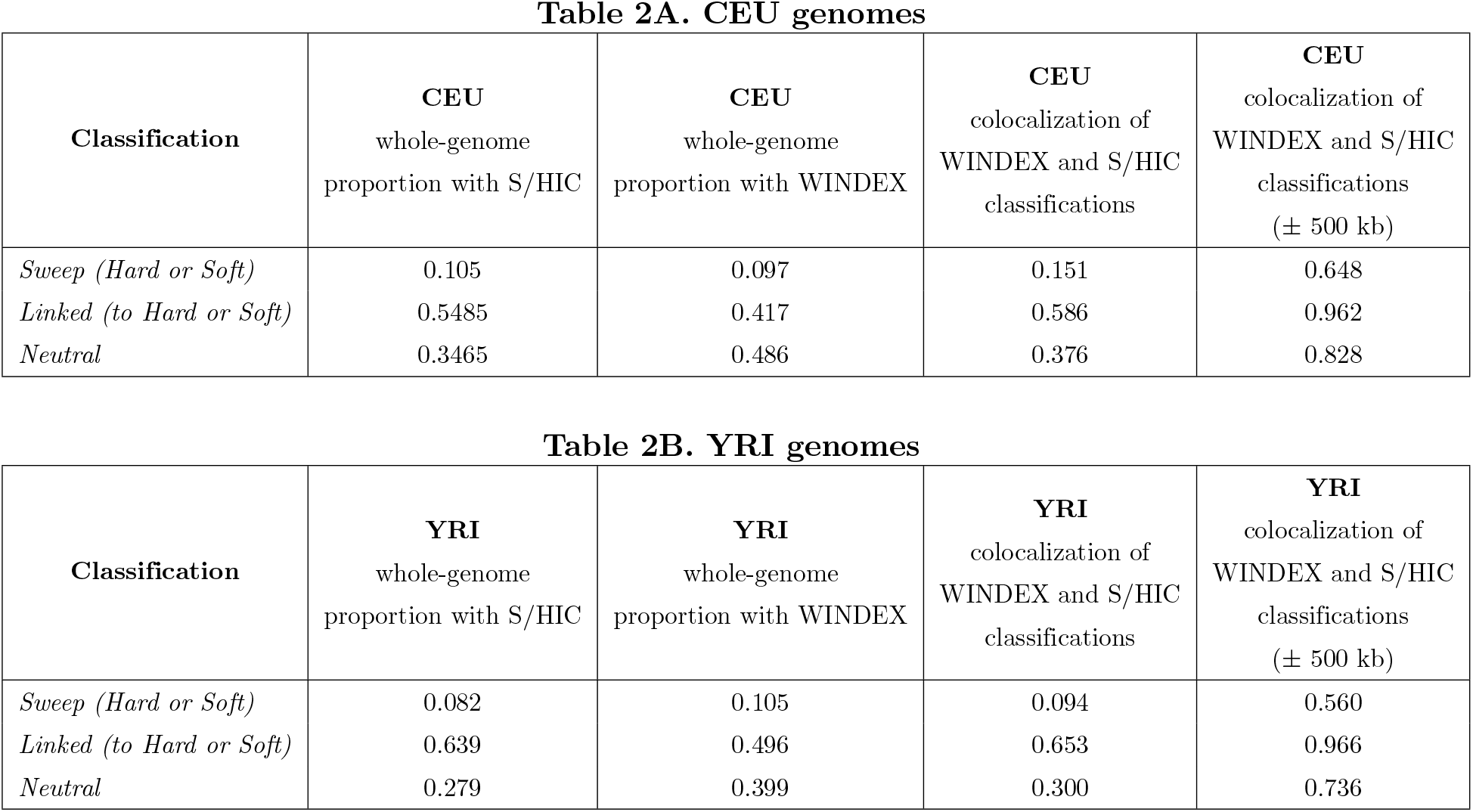
Comparative whole-genome scans with WINDEX and S/HIC reveal similar classification proportions of positive selection across populations.

We compared the classification outputs of WINDEX and S/HIC [16] in CEU and YRI genomes from 1000 Genomes Project data [18]. WINDEX only classifies based on three levels (“neutral”, “linked”, and “sweep”), so S/HIC results were releveled to match these classifications: “softLinked” and “hardLinked” were releveled to just “linked”, and “soft” and “hard” sweeps were releveled to just “sweep” (See Methods). In both CEU and YRI genomes, the whole-genome proportions of each class were calculated using both S/HIC and WINDEX across the same 100kb windows. Across both populations, WINDEX reports similar proportions of sweep regions and WINDEX reports consistently lower proportions of linked regions (columns 1 and 2 in both tables). Additionally, WINDEX classifications were colocalized with S/HIC classifications on both exact window match and within 500kb of the S/HIC prediction. When introducing this margin of error around matching classifications, the colocalization proportions across all three classes increased.

## Discussion

Here we present WINDEX, a hierarchical hidden Markov model that incorporates statistics at both the site-and window-based resolutions to localize targets of positive selection in regions of aligned population-genetic haplotype data. WINDEX can take into account specific demographic scenarios; by learning the distributions of selection statistics at linked sites in diverse populations, applications of WINDEX can offer new insights into demography-specific positive selection. Through the use of evolutionary simulations, we demonstrated that WINDEX outperforms our previous method, SWIF(r) [17], by leveraging inference from the site level to improve localization at the window-level, and increases specificity by using a restricted state structure within sweep windows (Figs. 2 and 3). Additionally, we compared the site-level performance of WINDEX with iSAFE [15], a selection statistic that ranks favored mutations that contribute to a positive selection region, in two canonical sites of positive selection in population samples from the 1000 Genomes Project. We observed that WINDEX was able to localize these loci well, and showed supporting evidence for these canonical sites when leveraged with a stochastic backtrace evaluation algorithm at the site level (Fig. 4). Finally, we compared the window-level classifications of S/HIC [16] and WINDEX across population whole-genome samples from the 1000 Genomes Project. These comparisons revealed very similar relative footprints of selection via classifications across both of these tools and showed moderately high colocalization of signals (Table 2).

There are multiple lines of future inquiry that could be further explored with WINDEX. First, a hierarchical version of the stochastic backtrace algorithm would provide uncertainty measures at the whole-genome level. This algorithm may better account for discordance between classifications from WINDEX and other tools, and would allow WINDEX to output more information for a user, as well as provide evidence for multiple putative loci in one region. Similarly, other post-hoc analyses could be applied to WINDEX for further LD interpretation, such as finite Markov chain imbedding (FMCI) and hybrid decoding [37]. Additionally, the emission framework in WINDEX is based off of a simple Naïve Bayes likelihood, and to more closely compare WINDEX with SWIF(r), WINDEX could be implemented with an emission framework that mimics the averaged one-dependence estimator underlying SWIF(r). This also may improve the ability of WINDEX to localize positive selection signals in more challenging scenarios, such as weaker selection. In our applications in this study, WINDEX has been trained only on strong *de novo* hard sweeps, but WINDEX can also be applied by training on other types of selection scenarios, such as balancing selection. Finally, more statistics could be used as input for training to better inform classification of other types of selection, such as the likelihood ratios T1 and T2 for balancing selection [38]. WINDEX is a powerful tool for understanding signals of positive selective sweeps in population-genetic haplotype data that offers new perspectives into the footprint of selection on genomes.

## Data and Code Availability

All necessary code to run WINDEX and related analyses can be found on our Github:

https://github.com/ramachandran-lab/WINDEX-lab/WINDEX

Real-world data used with WINDEX can be found at the 1000 Genomes Project:

https://www.internationalgenome.org/

## Acknowledgments

Thank you to David Peede for his help with polarization of the 1000 Genomes VCF files (Villanea and Peede *et al*. 2025). Code to replicate this is available at this Github repository:

https://github.com/David-Peede/MUC19-Peede/MUC19

